# Machine learning of honey bee olfactory behavior identifies repellent odorants in free flying bees in the field

**DOI:** 10.1101/2024.11.10.622876

**Authors:** Joel Kowalewski, Barbara Baer-Imhoof, Tom Guda, Matthew Luy, Payton DePalma, Boris Baer, Anandasankar Ray

## Abstract

Preventing beneficial insects like honey bees (*Apis mellifera*) from contacting pesticides on crops using odorants could counter current pollinator declines. However, the discovery of behaviorally aversive odorants is impeded by the complexity of the honey bee olfactory system where >180 olfactory receptors detect volatiles and generate valence. To solve this systems-level challenge we generated a machine-learning model to predict aversive valence from chemical structure using published olfactory behavior data in honey bees. We refine the predictive model by generating species level behavioral data for honey bees and *Drosophila* on an initial set of novel predicted repellents. The improved second computational model was then used to screen a chemical space of >50 million compounds and identify >130 repellent candidates. Behavioral validation using honey bees in the laboratory show a high predictive success. Additional testing of the top seven candidates using freely foraging honey bees in a field assay confirmed strong repellency, thus predicting a high probability to repel foraging bees from pesticide-treated crops. Machine learning, with iterative testing and modeling therefore provides a powerful approach for rational discovery of aversive volatiles for control of insects for which limited data is available.

**SIGNIFICANCE STATEMENT:** With honey bee populations declining partly due to pesticide exposure, we aimed to find smells that could keep bees away from pesticide-treated crops. We overcome challenges studying the complex bee olfactory system by developing an AI model trained on existing bee behavior data to predict chemicals bees would find aversive. The predictive model screened millions of compounds, identifying more than 130 potential repellents. Behavior testing in the lab and in field tests confirmed the effectiveness of the bee repellents. This method could lead to bee-safe pesticide formulations, potentially protecting pollinator populations while maintaining crop protection.

## MAIN TEXT

Honey bees play a critical role as pollinators for nearly a third of food crops (*1*). However, over the last decade, beekeepers reported increasingly unsustainable losses (>60%) of their managed hives every year. Several factors have been identified contributing to these declines, amongst them parasites such as Varroa mites, climate change, and pesticides (*2–4*).

When honey bee workers forage in pesticide-treated fields, they bring back pesticide residues into their hive, which negatively impacts the survival of their colony, endangering the pollination of high-value crops, even at sublethal dosages. We propose a novel solution for this problem: identifying honey bee repellents that can be added to pesticides, thus reducing their contact with treated crops. Honey bees have a powerful sense of smell and use their antennae to detect numerous volatile chemicals which they sense using a sophisticated olfactory system of >160 Odorant receptor genes, and 21 IR genes in the genome which encode for odor-gated ion channels (*5, 6,*). However, the complexity of the insect olfactory system makes it a significant challenge to experimentally identify repellent odorants. In fact, in the 20 years since the odorant receptor families in insects have been identified, not a single new repellent odorant was registered for use in a new insect repellent product by the U.S. Environmental Protection Agency.

Recently we used Machine Learning to successfully create structure-based models to predict mosquito repellents from chemical structure (*7*) and for human olfactory behavior allowing us to predict ∼140 odor characters with great success (*8, 9*). While a large empirical dataset of repellent odorants is available for mosquitoes, comparable data for honey bees are much more limited. (*7*). Here we used a related Machine Learning approach, scanning 3D molecular structures of odorants *in silico,* to predict their potential for repelling honey bees (Figure 1). We overcame the challenges of a small set of training odorants by performing a second round of modelling using data from an initial round of testing to improve the predictive ability (Figure 3A).

**Figure 1.**
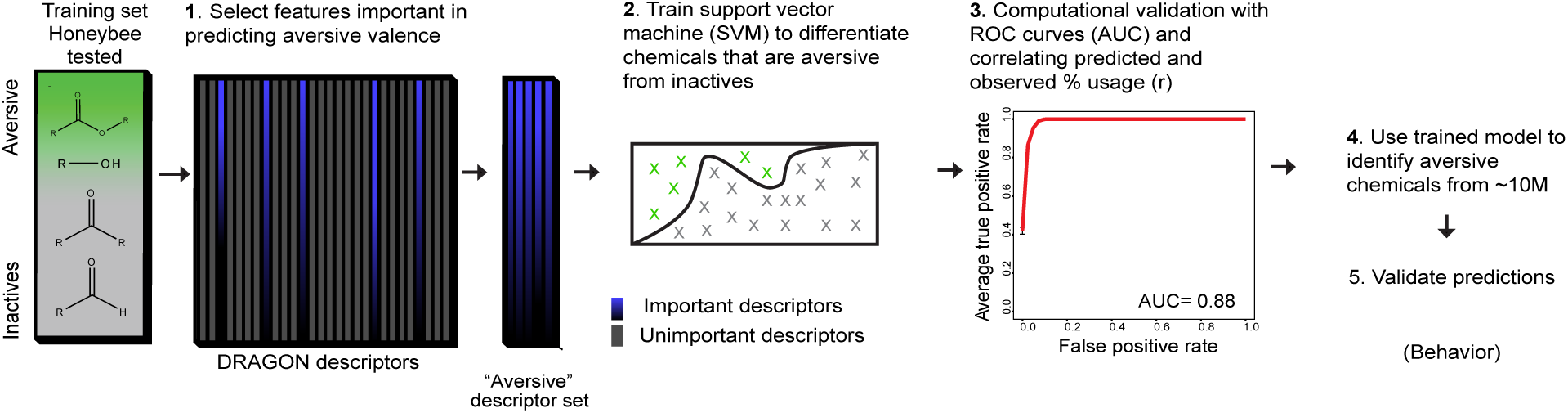
A chemical informatics method to predict honey bee repellence. Overview of the steps for the Machine Learning-based cheminformatics pipeline used to model honey bee repellence from the 3D structure of a training set of known repellents, and subsequent in-silico screening for novel repellents from a large chemical space.

To do so, we first reviewed the existing literature on honey bee olfactory behavior and assembled a training set of volatile chemicals including a subset with a negative behavioral valence (*10–14*). We generated energy-minimized 3D structures for each chemical and calculated 5,290 physicochemical features: initially for 184 training chemicals and later for 203 chemicals. From the initial training set we identified 45 features that were considered important for predicting aversive valence (Table S1). We then evaluated a collection of Machine Learning (ML) algorithms including regularized Random Forest, Gradient Boosted Decision Trees, and a non-linear Support Vector Machine (e.g., incorporating the Radial Basis Function kernel) with these features to predict repellency of the test chemicals To cross-validate each algorithm’s performance, we divided the training data into multiple smaller training data sets and test data sets and calculated average predictive success, graphically depicted as the area under the Receiver Operating Characteristic curve (ROC AUC) in our model. Our computational validation (avg AUC = 0.88) indicated that our model was able to distinguish volatiles with aversive valence from non-repellents (Figure 1).

Using this model, we subsequently evaluated a library of ∼45M small molecules from the MolPort Database. The top predicted repellents (∼7% of hits with computed repellency scores >=0.9), represented a structurally diverse group of 139 chemicals, of which some showed similarities to the known repellents in the training set. The notion that Machine Learning can predict the behavioral valence of small molecules towards honey bees from just 3D chemical structure would be revolutionary and inform follow up behavioral validation.

Because our training set included a few known repellents for other insects as well, such as benzaldehyde and geraniol, we anticipated that the predictions likely included broad-spectrum insect repellents alongside honey bee-selective ones. If repellents were to be mixed with insecticides to repel bees, it might be desirable not to repel pest insects. To evaluate predicted compounds for broad-spectrum insect repellent activity we performed behavior assays using another insect, the fruit fly *Drosophila melanogaster* (Figure 2A). We randomly selected a set of compounds from the predictions and used a trap-based assay to quantify fly aversion. Several compounds showed low levels of repellency, with a <50% reduction in numbers of flies entering traps relative to the solvent control (Figure 2A). Subsequently, we only used those compounds for further tests in honey bees.

**Figure 2.**
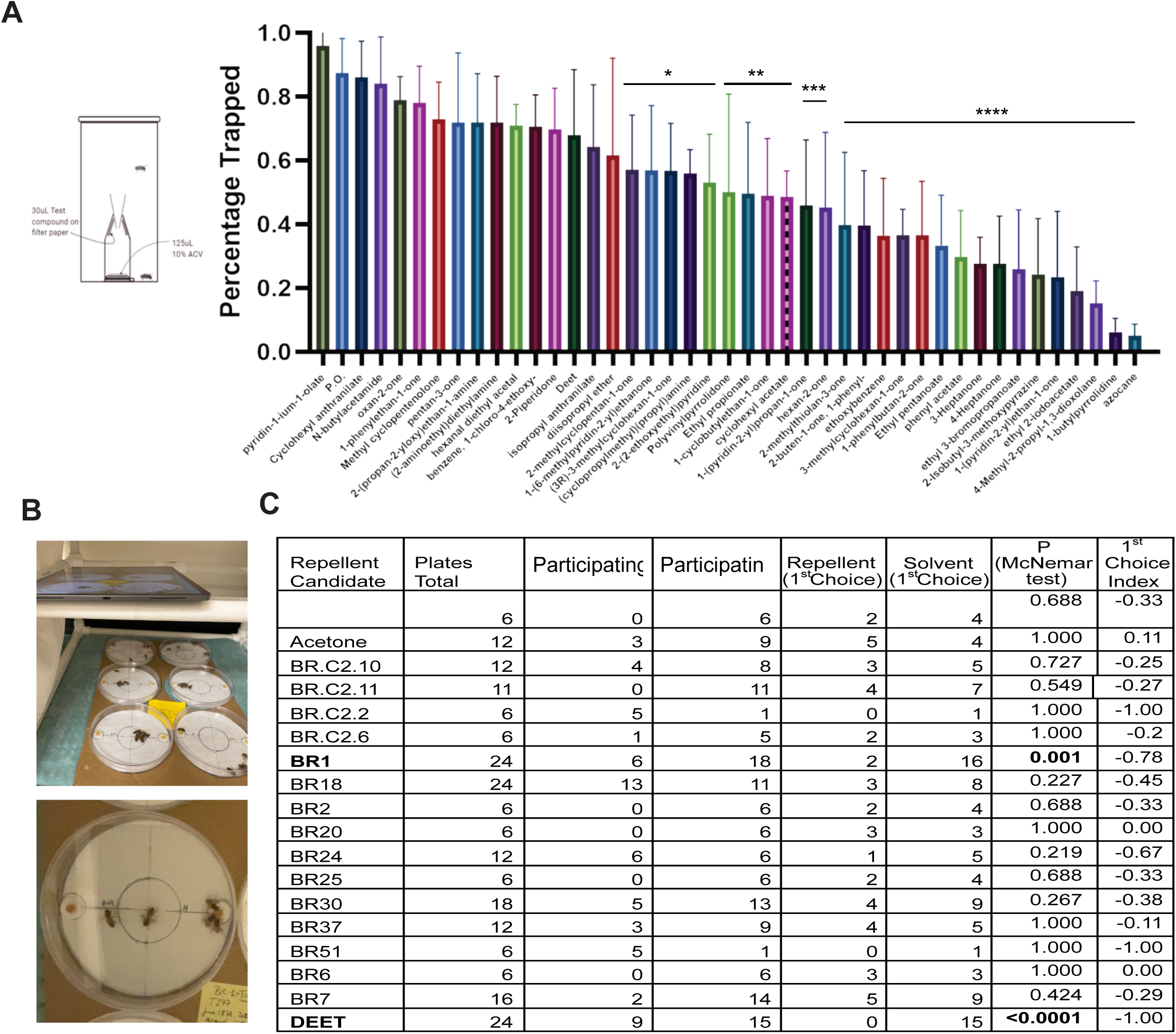
Predicted honey bee repellents. **A.** Testing Chamber containing a one-choice trap to determine, whether an odorant will repel fruit flies, and the mean percentage of fruit flies caught in a trap treated with predicted repellent odorants (10% v/v in Paraffin oil) and baited with 10% apple cider vinegar. N=5-8 trials (∼20 flies/trial). Error bars= s.e.m. * p<0.05, ** p<0.005, *** p<0.001, **** p<0.0001. **B.** Photograph of the two-choice Petri dish arenas used to test the behavior of honey bees to repellent candidates individually, and **C.** Table with preference indexes in the two-choice assay for the first choices of honey bee workers offered two containers of honey water (50 μl of 50%) placed on filter paper discs treated with a test chemical or treated with solvent alone. The index was calculated for each compound as (total number of repellent choices minus total number of solvent choices) divided by the sum of all tests. We used McNemar tests for pairwise comparisons.

We modified a honey bee repellent testing assay in Petri dishes in the lab to test the 15 candidate compounds (*15*). We placed honey-water (50μl) filled microfuge lids onto treated filter paper disks on two sides of a Petri dish and placed ∼five hungry honey bee workers in the center. From video recordings we evaluated whether the first bee navigated to a honey pot on the repellent treated or the control solvent side (Figure 2B). For many of the tested compounds the bees preferred to visit the honey-water pots on the control side versus the repellent side, (Figure 2C).

The initial training data set used for Machine Learning was limited in size and lacked high quality quantitative values for structurally related compounds. The predictive models can be improved substantially by adding new compounds with quantitative data that are structurally related to the training set. We therefore developed an iterative pipeline where the previous round of behavioral validation data is added to the training data and used to rebuild a more refined training set (Figure 3A). Specifically, we added the quantitative data from the 15 newly tested compounds to the training set. The 3D structures were analyzed, and a physicochemical feature set was selected as before (Table S1). As expected, the computational validation results based on the AUC values, show an improvement. We next screened a ∼10M compound library in silico with the updated repellent predictive ML model and identified a wide array of natural and synthetic honey bee repellent candidates (Figure 3A). Because the rate of evaporation may affect how long a repellent will be effective in the field, we estimated the vapor pressure for each candidate. To do this, we used Machine Learning on a newly created training data set of 1,931 compounds with known vapor pressures.

**Figure 3.**
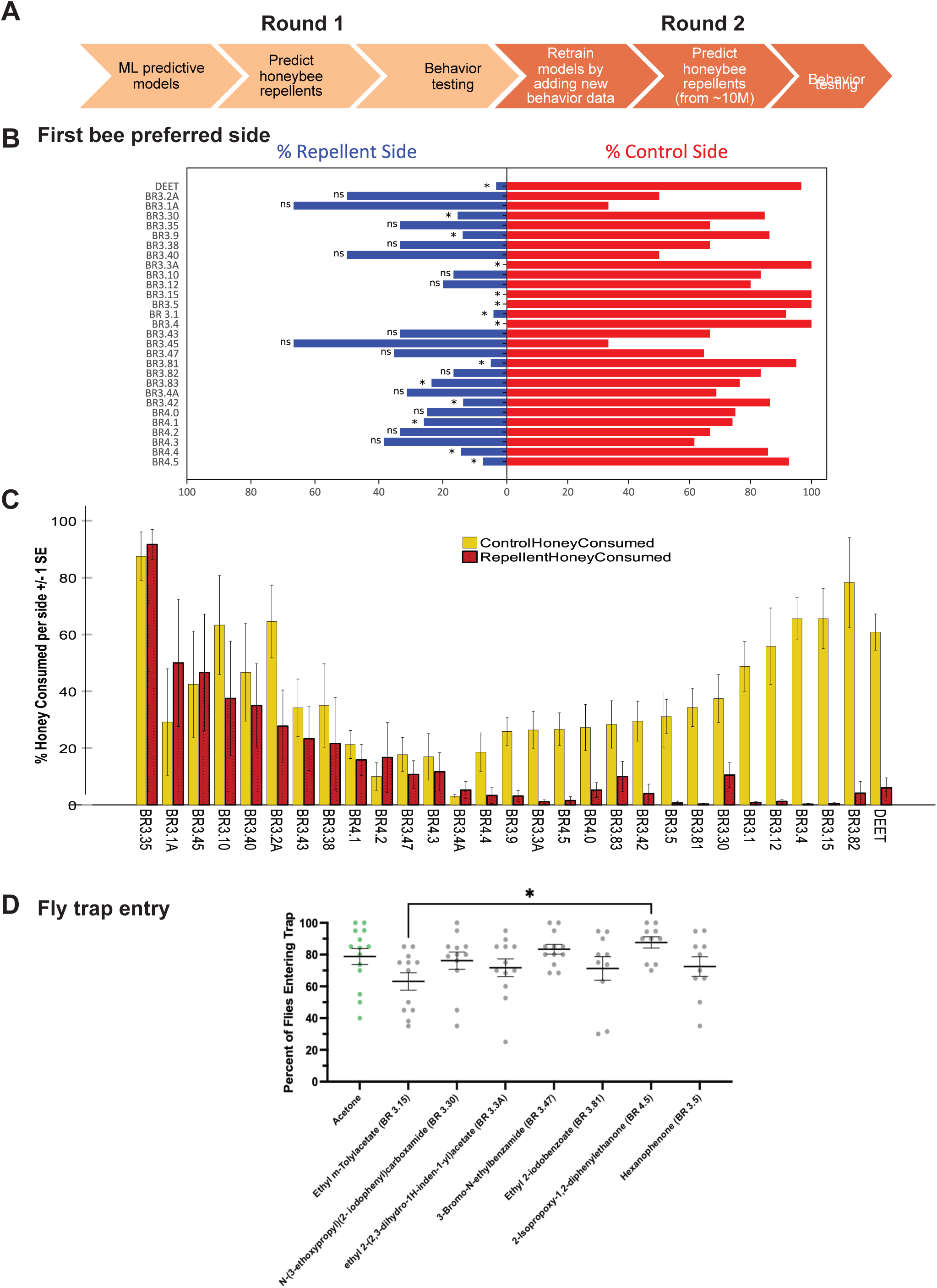
Reiterative training of ML predictive models and testing honey bee repellents. **A.** Schematic pipeline for the overall computational-behavior hybrid pipeline showing the two rounds. **B.** Mean Preference index of honey bees in making the first choice to move to the repellent treated side in a two-choice plate assay. 70 μl of pure honey was used in the two containers to attract the bees. **C.** Average consumption of honey from containers placed on each side in the two-choice plate assays (N = 460 plates). The index is calculated for each compound as which honey pot has greater consumption (total number of repellent side with greater consumption minus total number of solvent side with greater consumption) divided by sum of all choices). **D.** Mean percentage of fruit flies caught in a trap treated with predicted repellent odorants at application rates to test in field tests (0.1mg/cm^2^) and baited with 10% apple cider vinegar. N=5-8 trials (∼20 flies/trial). Error bars= s.e.m.

To validate the second round of predictions and identify compounds with the strongest behavioral responses in honey bees, we selected 28 molecules with high predicted repellency values that fell into one of three vapor pressure (V.P.) categories: very low V.P solids, very low V.P liquids (10^-4^ - 10^-3^ mmHg @25C) and low V.P. liquids (10^-2^ mmHg @25C). We tested the compounds on honey bee behavior using the 2-choice arena as before but used pure honey (70μl) instead of honey water. Using honey increased the bees’ participation from 68% in round 1 to 81% in round two, and the increased volume allowed us to evaluate honey consumption on each side of the arena after one hour. Most of the predicted repellent compounds showed strong repellency (Figures 3B). The repellent effects were conserved across three or more different honey bee colonies when tested in batches demonstrating a robust and reproducible effect (Figure S1). For 15 of the 28 test compounds, we found that the bees drank significantly less honey on the treatment side compared to the control side as well (Figure 3C, Table S2).

We next tested seven compounds that showed the strongest repellency in honey bees on *Drosophila melanogaster* using the 24-hr trap assay as before, at a concentration relevant for field applications as done later (∼0.1 mg/cm^2^). The flies showed no behavioral aversion to the candidate repellent chemicals, compared to controls (Figure 3G). This suggests that the new honey bee repellents were not aversive to other insects like fruit flies.

Next, we tested the seven repellents on freely flying foraging honey bees in a field assay. Honey bees are readily attracted to wax honeycomb foundations that have been sprayed with sugar-water. We sprayed sugar-water on three equally spaced wax-foundations placed near hives and treated them with either (1) solvent (acetone), (2) a candidate honey bee repellent (rate of ∼0.1 mg/cm^2^) or (3) a positive control (DEET) at the same rate (Figure 4A). The honey bees flew to the wax foundations and were offered a choice between the different treatments. We filmed the assays and counted the bees on each foundation in five-minute increments. Each of the seven compounds showed repellency to honey bees, with three of them showing aversion up until the end of the assay. We ended the assay counts when the honey bees stopped being attracted to the untreated controls (Figures 4A,B,C). Kruskal-Wallis tests revealed significantly lower numbers of bees on treated wax foundations compared to controls (Figure 4C), In addition, we tested the two top-scoring compounds at half their original concentration (0.05 mg/cm^2^) and still observed significant repellency (Figure 4D).

**Figure 4.**
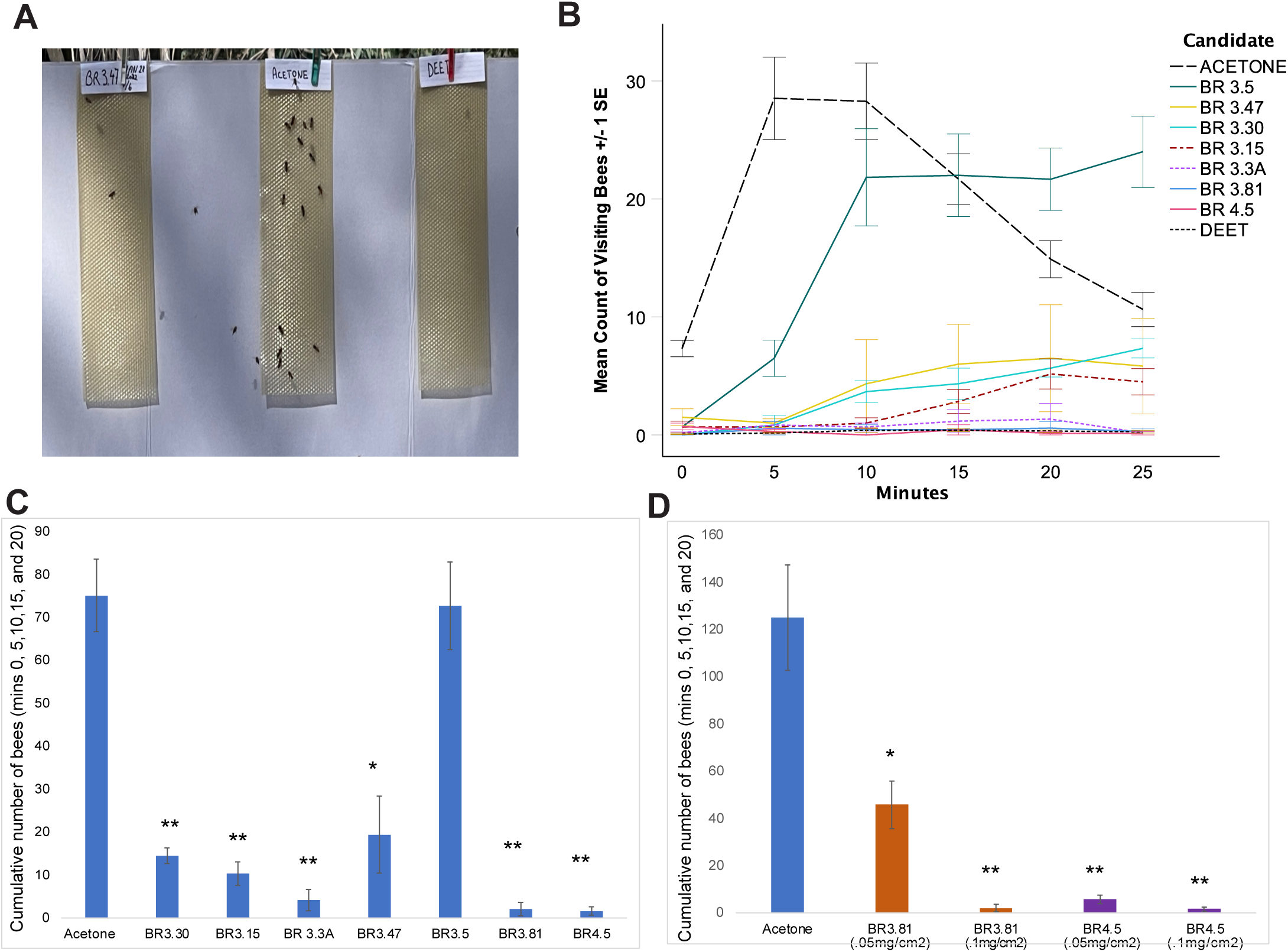
Field testing of top repellents in the lab using robbing assays. **A.** Representative photo of a robbing assay. **B.** Time-course of the mean numbers of foraging honey bees visiting for each indicated compound tested (0.1 mg/cm^2^). DEET was used as a positive control and acetone as a negative control in each assay. N=6-7. Error bars=s.e.m. **C.** Mean number of honey bees for each compound summed over the time points shown in **B.** * p<0.05, ** p<0.005, *** p<0.001. We used Kruskal Wallis tests for pairwise comparisons. **D.** Mean number of honey bees for each compound tested at the regular dose (0.1mg/cm^2^) and half dose (0.05mg/cm^2^), summed over the time points over the duration of the experiment as above. N=6, Error bars=1 s.e.m.

Taken together our results from the lab and field assays demonstrate that a computational approach can successfully model honey bee aversive valence from chemical structure alone and can be used to efficiently screen millions of chemicals to find strong repellents.

## Discussion

Most odorants reaching a bee’s antennae activate or inhibit several of the >160 olfactory receptors at the same time. Each odorant-receptor interaction elicits a specific neuronal activity in the bee brain, consequently allowing for a limited number of receptors to encode a practically infinite number of olfactory combinations, with >160 OR and 21 IR genes in the genome (reviewed in(*16*)). Even though honey bees are a well-studied model system for olfactory coding and learning (*16*), (*17*), we still know little about the pathways between olfactory stimulus evaluation and decision-making (*16*). The olfactory system therefore poses an extreme challenge to modeling. In spite of what is already known about the honey bee olfactory system, it would take several years and significant funding before generating a near complete knowledge of the olfactory system that allows to do predictive modelling of valence. The 3D structure based computational approach fills a very important void in identifying novel volatiles triggering a specific olfactory behavioral response in honey bees. This machine learning pipeline can accelerate research into insect repellency and its physicochemical basis. The overall species-selective approach we have presented will help identify additional novel repellent chemicals for honey bees as well as inspire an exploration of the vastness of chemical space for use in repellent strategies for other animals.

Pesticides have been identified as a significant contributor to the global decline in honey bee populations (*18, 19*). Numerous studies have demonstrated the detrimental effects of various pesticides alone or in combination with other environmental stressors on honey bee health, including neonicotinoids, organophosphates, pyrethroids (*20, 21*) and bifenazate (*22*). These chemicals can cause acute toxicity, leading to immediate bee mortality, as well as sublethal effects that impair bee behavior, navigation, immune function and population growth (*22, 23*). Chronic exposure to pesticides, such as imidacloprid, has been linked to reduced colony growth, queen failure, and increased susceptibility to pathogens and parasites (*24*). The widespread use of pesticides in agriculture has raised concerns about the sustainability of honey bee populations and the potential impacts on pollination services and food security (*19*).

Recent research has explored the potential of using bee repellent odors to mitigate the harmful effects of pesticides on honey bees. The concept involves applying non-toxic, natural compounds that emit odors that are unappealing to bees, thereby deterring them from foraging in treated areas during pesticide application. By reducing bee exposure to pesticides, this approach aims to minimize the direct and indirect impacts on bee health. Studies have identified several promising bee repellent compounds, such as 2-heptanone and essential oils from plants like lavender and thyme but their low efficacy does not make them practical in the agricultural setting (*25, 26*). Our field behavior assays strongly suggest that the use of bee repellent odors can significantly reduce bee activity in treated areas, potentially lowering the risk of pesticide exposure. This study also provides a foundation for further research to optimize the effectiveness of bee repellent odors, assess their long-term impacts on bee behavior and colony health, and develop practical application strategies for integration into pest management practices. Moreover, our approach for a rational and high-throughput discovery of bee repellents can be extended to any other species of interest, thus providing a very powerful and affordable computational tool for chemical ecologists.

## MATERIALS AND METHODS

### Machine learning repellency

Each chemical structure tested for attraction/repellency on bees was prepared for machine learning (ML) using RDKit in Python. The 2D structures were first embedded in 3D coordinate space; the 3D geometries were then optimized using the Merck molecular force field (MMFF94). For each optimized 3D structure, AlvaDesc computed 5,290 diverse 2D and 3D chemical features, describing large-scale properties such as molecular shape that affect sensory receptor docking in addition to forces describing potentially more abstract interactions leading to graded activation or inhibition of the sensory receptors of the bee.

Bee behavioral data, presented as a preference score (5 = max repellency), was converted into a binary label, repel/inactive, using > 1 as the cutoff between repellent and neutral or attractant behaviors. In other circumstances, bee behavior to compounds was reported without a repellency score. For attractive compounds, an inactive label was assigned based on previous data (*11–14, 27, 28*). The data were subsequently split into training and testing partitions (90:10) and any further processing was done exclusively on the training partition.

The large chemical feature matrix was next reduced to a subset of features that optimally classified compounds according to their repellent activity on bees. This involved using the Recursive Feature Elimination (RFE) algorithm alongside a Support Vector Machine (SVM) with a Radial Basis Function (RBF) kernel to evaluate the feature importance over 250 partitions of the training data (e.g., 10-fold cross validation, repeated 25 times). An intermediate subset >100 features was next passed to a series of algorithms (e.g., Random Forest and Boosted Decision Trees) to re-rank the features, selecting the final, consensus set.

Top features were evaluated across a diverse set of algorithms including SVMs, Decision Trees, and Neural Networks (Multilayer Perceptron), and the cross validated classification metrics suggested high classification success rates (AUC, Sensitivity, and Specificity) when aggregating the predictions of three algorithms: SVM with a Radial Basis Function (RBF) kernel, a Gradient Boosting Machine (built with Decision Trees), Random Forest with regularization. Final Evaluation was done on the test set. Trained models predicted 60,000 metabolites and other naturally sourced compounds. The MolPort database, which contains 47 million compounds in total, was filtered to a smaller subset for repellency predictions, based on commercial availability and structural similarity to known bee repellents in the training set. After the first round of experimental validation tests with bees, the models were retrained. Additional prospective repellent compounds were scored using the updated models.

### Machine learning Vapor pressure

Training and testing data were obtained from EPI Suites, Environmental Protection Agency (EPA). Methods for fitting these models are as outlined for bee repellency; namely, chemical features were computed using AlvaDesc, followed by processing the feature values and identifying a small predictive subset through the recursive feature elimination algorithm (RFE). To compare the vapor pressure model predictions with respect to different machine learning methods as well as EPI suite, data were split according to the train/test partitions defined in a previous study using the vapor pressure data for Machine Learning (ML) (*29*).

### Fly trap assay

*Drosophila melanogaster* stocks (RRID:BDSC_1) maintained in the lab were grown in standard media bottles. All chemicals were purchased at highest purity through Molport. Flies of a similar age (4-6 days) were collected in groups of 20 (10 females +10 males), and wet-starved for 24hrs in starvation vials. Testing chambers (Dram vials with centrifuge tube traps containing lure (10% Apple Cider Vinegar) plus test compound of desired dilutions) were assembled and flies quickly transferred to them by gentle tapping. The assays were left on the bench and data on number of flies entering traps recorded after18 hours.

### 2-choice honey bee plate assays

Honey bee (*Apis mellifera*) workers for our experiments were bred in apiaries at the University of California Riverside, between August 2021 and April 2022. To age them, we placed a frame of capped worker brood at the red-eye stage into an observation frame in a humidified incubator at 34°C. After two to three days, we collected groups of 80 newly hatched workers into specially 3D-printed experimental cages (12×10×7.5cm), where we kept them until they were reached an age of 13 to 19 days, corresponding to foraging age). The bees received sugar water (50%) and water ad libitum, as well as a paste of protein powder (https://megabee.com/) mixed with 50% sugar water every second day until day eight.

The behavior arenas consisted of plastic Petri dishes with a diameter of 15 cm with two filter paper circles on each side (1 cm diameter) treated with 20 μl of solvent (acetone evaporated for 30 min) on one side, and 20 μl of test compound (20 μl, 5% in acetone, evaporated for 30 min) on the opposite side of the arena. To stimulate bees to participate, 50 μl of 50% honey-water in cut caps of PCR tubes was carefully taped to the center each of the two treated papers (Figure 2). For round two (Figure 3), 70 μl of pure (slightly warmed) honey was used in each cap (the honey surface formed an upward meniscus). The night before the experiment, the bees were water-starved, then cooled down at 4°C for approximately 20 minutes before placing five bees into the middle of the arena. The spacing of the filter papers was kept consistent by taping a template (round piece of paper) to the bottom of each Petri dish and taping the filter papers on top of the correct spot onto the template. An Apple iPad or iPhone was used to film six arenas at once for one hour in a custom-built PVC tube cage covered with a cotton sheet to create a uniform visual surrounding. To wake up the bees, the choice arenas were placed onto a heating pad (20’’x24’’, Model No PEHPWIDE-SG, Pure Enrichment, purenrichment.com, set to level 1).

Videos were analyzed offline. For each plate, the side the first bee chose to drink honey from first was recorded (first choice). An additional semi-quantitative metric was noted for Figure 3: an estimate of the percentage of honey consumed in each container 95% (meniscus gone, only), 75%, 50%, 25%, or 0%. Plates in which the bees had not consumed any honey from either side (non-participating plates) were excluded from further evaluations. For each batch of six plates, workers from the same colony were used. For repellent candidates with the most promise, bees from at least three different colonies were tested (at least six participating plates each) (Table S2).

### Robbing assays in the field

For this assay we elicited the robbing behavior of honey bees, which means that they steal unguarded honey or honey guarded by weaker colonies (*30*). To do so, we sprayed standardized wax foundations (44 x 25.5 cm) with 50 puffs of 50% honey diluted in water as an attractant. Chemicals were sprayed on top at 0.1mg/cm^2^ (15 puffs of 5% in acetone). Three foundations, solvent, DEET and test compound, were affixed on a standing poster board (92×122 cm) placed on a table five meters in front of the first row of 40 hives and filmed for robbing behavior by the bees for 30 min, using an iPad or an iPhone. Prior to the start of the days experiments, three honey frames and honey lures were placed on a table for ∼15-30 min to trigger robbing behavior of bees from nearby colonies and then removed before trials. The position of the three wax foundations to each other was randomly shifted. After 25 min, the positive control sugar-water sprayed wax foundations were no longer attractive to honey bees and the analysis was restricted to this time point. We repeated each assay a minimum of six times. Moreover, we repeated the assays with the two most promising candidates at half the first concentration (∼0.05 mg/cm^2^) four times each.

### Statistics

To compare the proportion of plates per test batch in which the bees chose to drink honey from the control side versus the treated sides, we used non-parametric, pairwise McNemar tests, assuming the null hypothesis of an equal distribution between choices or 50/50. The non-parametric, pairwise Wilcoxon Signed-Rank tests were used to compare the percentage of honey consumption between treatments. Plates in which bees did not drink any honey were removed (non-participating plates). For the assays on Drosophila behavior, we performed ANOVA followed by Tukey’s multiple comparison tests to compare the number of flies entering the trap between treatments and control. For the robbing assays, we summed up the counts of visiting bees on each wax foundation at 5, 10, 15 and 20 minutes, then used non-parametric Kruskal Wallis tests for pairwise comparisons between each of the three treatments. We adjusted the p-values for multiple testing with Bonferroni corrections.

## RESOURCE AVAILABILITY

### Lead Contact

Anandasankar Ray

### Materials Availability

No new materials were generated in this study.

### Data and Code Availability

Data used in the analyses is publicly available from the references cited, and the data generated in this manuscript are supplied within the figures. Any additional data associated with this manuscript is also available in other formats on request from the corresponding author and lead contact. The code developed for this study is available from the corresponding author for non-commercial use upon request. Use of the code may be subject to a standard Material Transfer Agreement (MTA) or specialized licensing agreement through University of California Riverside’s Technology Transfer Office, as patent applications related to this software are currently pending.

## ACKNOWLEDGEMENTS

This work was supported by funding provided by the California Research Alliance (CARA) by BASF. The content is solely the responsibility of the authors and does not represent the official views of the funding agencies. The funders had no role in study design, data collection and analysis, decision to publish, or preparation of the manuscript.

## FIGURE LEGENDS

**Figure S1.**
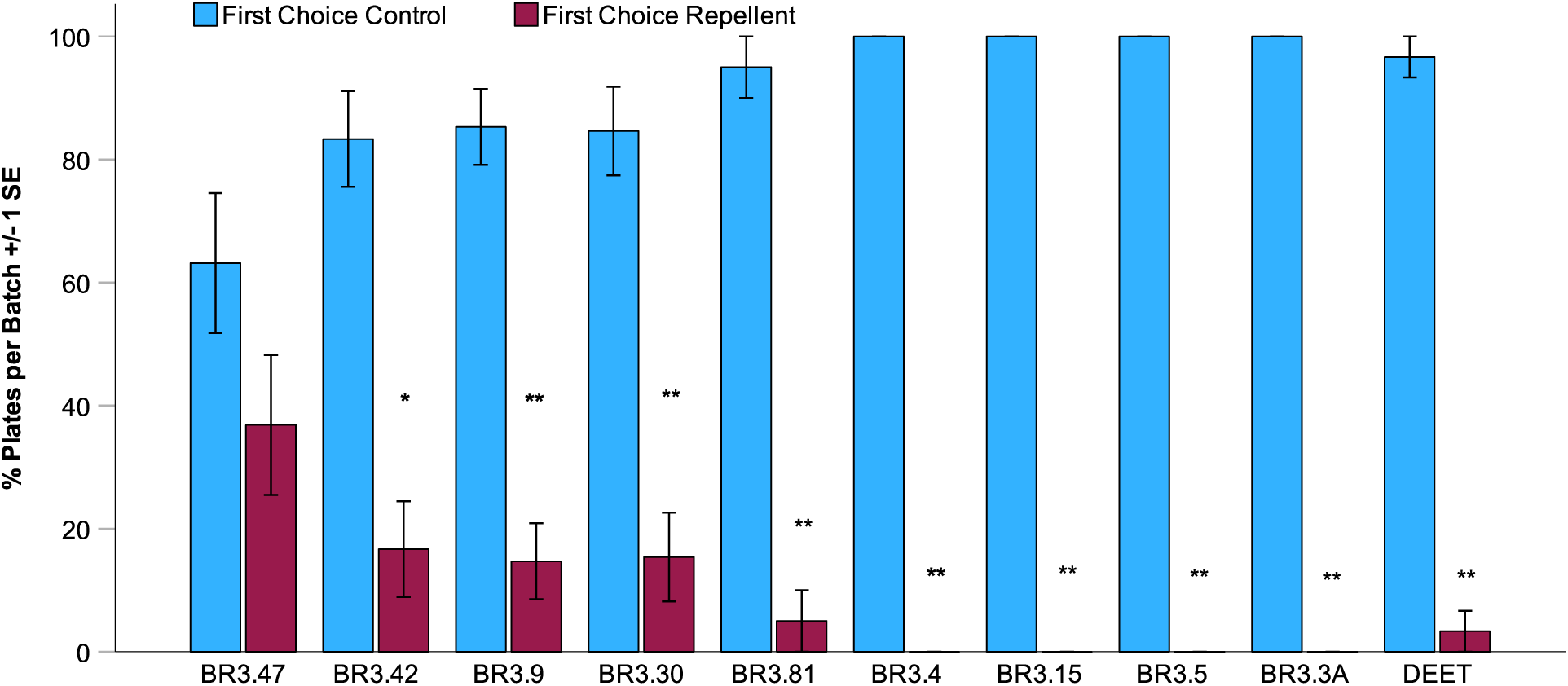
**(related to Figure 3)**. Average percentage of two-choice plate assays in which honey bees made their first choice to either move to the repellent treated side or solvent treated side across different honey bee colonies, N=3-6 colonies (∼6 plates/colony). Error bars =1 s.e.m., P*<0.05, **<0.001.

**Table S1.** Optimized DRAGON physicochemical descriptor set from iterative training in Figure 3A used to predict bee repellent compounds.

| Name | Description |
| --- | --- |
| Mor28e | signal 28 / weighted by Sanderson electronegativity |
| CATS2D_03_NL | CATS2D Negative-Lipophilic at lag 03 |
| Mor28s | signal 28 / weighted by I-state |
| Gu | total symmetry index / unweighted |
| Eta_B | eta branching index |
| RDF040m | Radial Distribution Function - 040 / weighted by mass |
| ATSC4m | Centred Broto-Moreau autocorrelation of lag 4 weighted by mass |
| RDF040v | Radial Distribution Function - 040 / weighted by van der Waals volume |
| DLS_03 | modified drug-like score from Walters et al. (6 rules) |
| GATS8i | Geary autocorrelation of lag 8 weighted by ionization potential |
| DISPe | displacement value / weighted by Sanderson electronegativity |
| Mor17m | signal 17 / weighted by mass |
| MAXDN | maximal electrotopological negative variation |
| TDB01e | 3D Topological distance based descriptors - lag 1 weighted by Sanderson electronegativity |
| RDF035p | Radial Distribution Function - 035 / weighted by polarizability |
| GATS2s | Geary autocorrelation of lag 2 weighted by I-state |
| VE3sign_Dz(v) | logarithmic coefficient sum of the last eigenvector from Barysz matrix weighted by van der Waals volume |
| H2s | H autocorrelation of lag 2 / weighted by I-state |
| E1s | 1st component accessibility directional WHIM index / weighted by I-state |
| SpDiam_AEA(dm) | spectral diameter from augmented edge adjacency mat. weighted by dipole moment |
| ATSC2s | Centred Broto-Moreau autocorrelation of lag 2 weighted by I-state |
| X5Av | average valence connectivity index of order 5 |
| Mor27s | signal 27 / weighted by I-state |
| P2m | 2nd component shape directional WHIM index / weighted by mass |
| SpMax2_Bh(s) | largest eigenvalue n. 2 of Burden matrix weighted by I-state |
| P_VSA_p_2 | P_VSA-like on polarizability, bin 2 |
| IVDE | mean information content on the vertex degree equality |
| L/Bw | length-to-breadth ratio by WHIM |
| TDB03m | 3D Topological distance based descriptors - lag 3 weighted by mass |
| SM15_EA(ri) | spectral moment of order 15 from edge adjacency mat. weighted by resonance integral |
| H4p | H autocorrelation of lag 4 / weighted by polarizability |
| CATS2D_04_NL | CATS2D Negative-Lipophilic at lag 04 |
| VE2sign_Dz(v) | average coefficient of the last eigenvector from Barysz matrix weighted by van der Waals volume |
| GGI1 | topological charge index of order 1 |
| Mor28m | signal 28 / weighted by mass |
| RTs+ | R maximal index / weighted by I-state |
| Mor21s | signal 21 / weighted by I-state |
| SpMAD_X | spectral mean absolute deviation from chi matrix |
| RDF035i | Radial Distribution Function - 035 / weighted by ionization potential |
| Mor28v | signal 28 / weighted by van der Waals volume |
| MATS1e | Moran autocorrelation of lag 1 weighted by Sanderson electronegativity |
| RDF010s | Radial Distribution Function - 010 / weighted by I-state |
| TDB06s | 3D Topological distance based descriptors - lag 6 weighted by I-state |
| P_VSA_LogP_3 | P_VSA-like on LogP, bin 3 |
| VE2sign_G/D | average coefficient of the last eigenvector from distance/distance matrix |

**Table S2.**
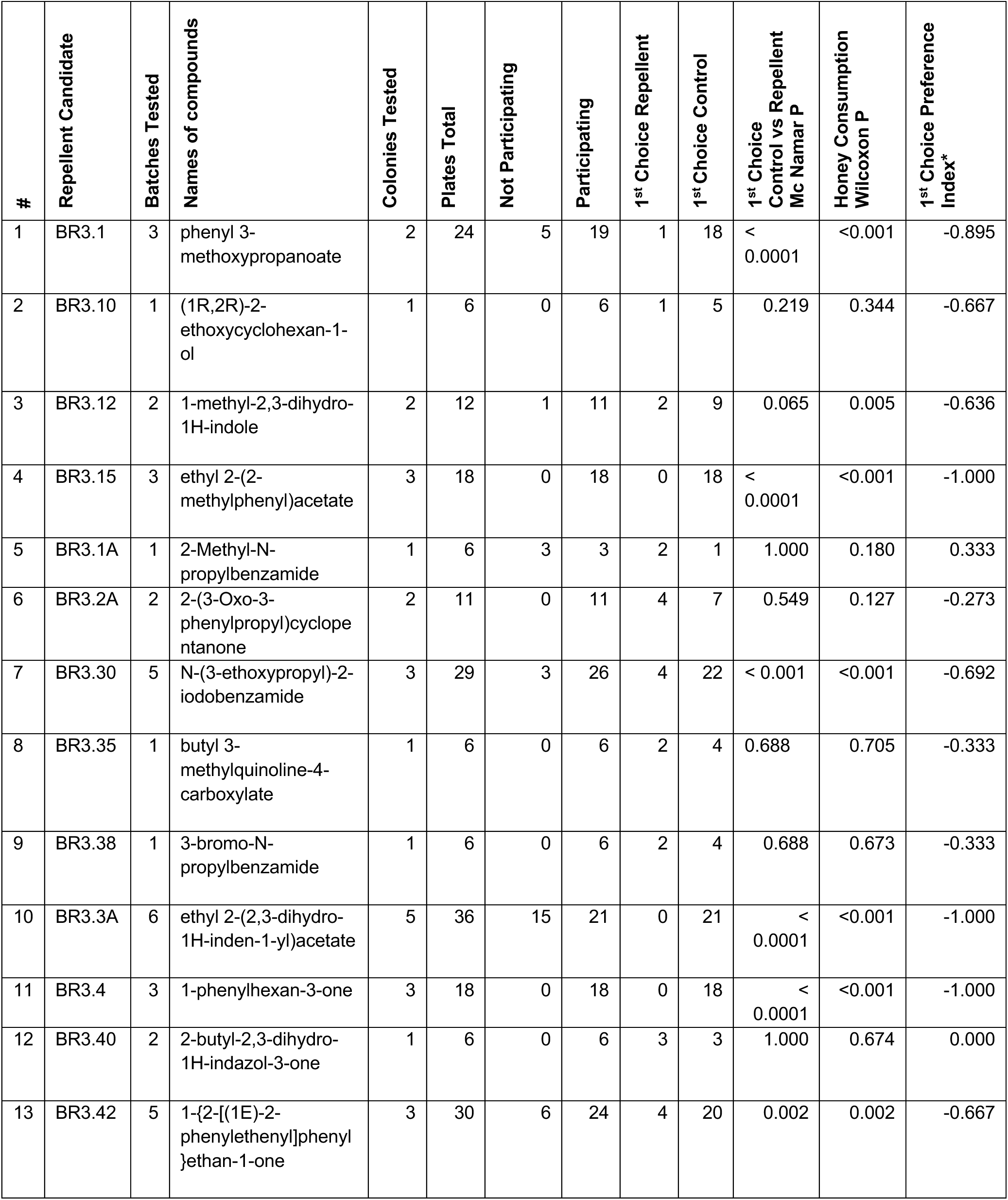
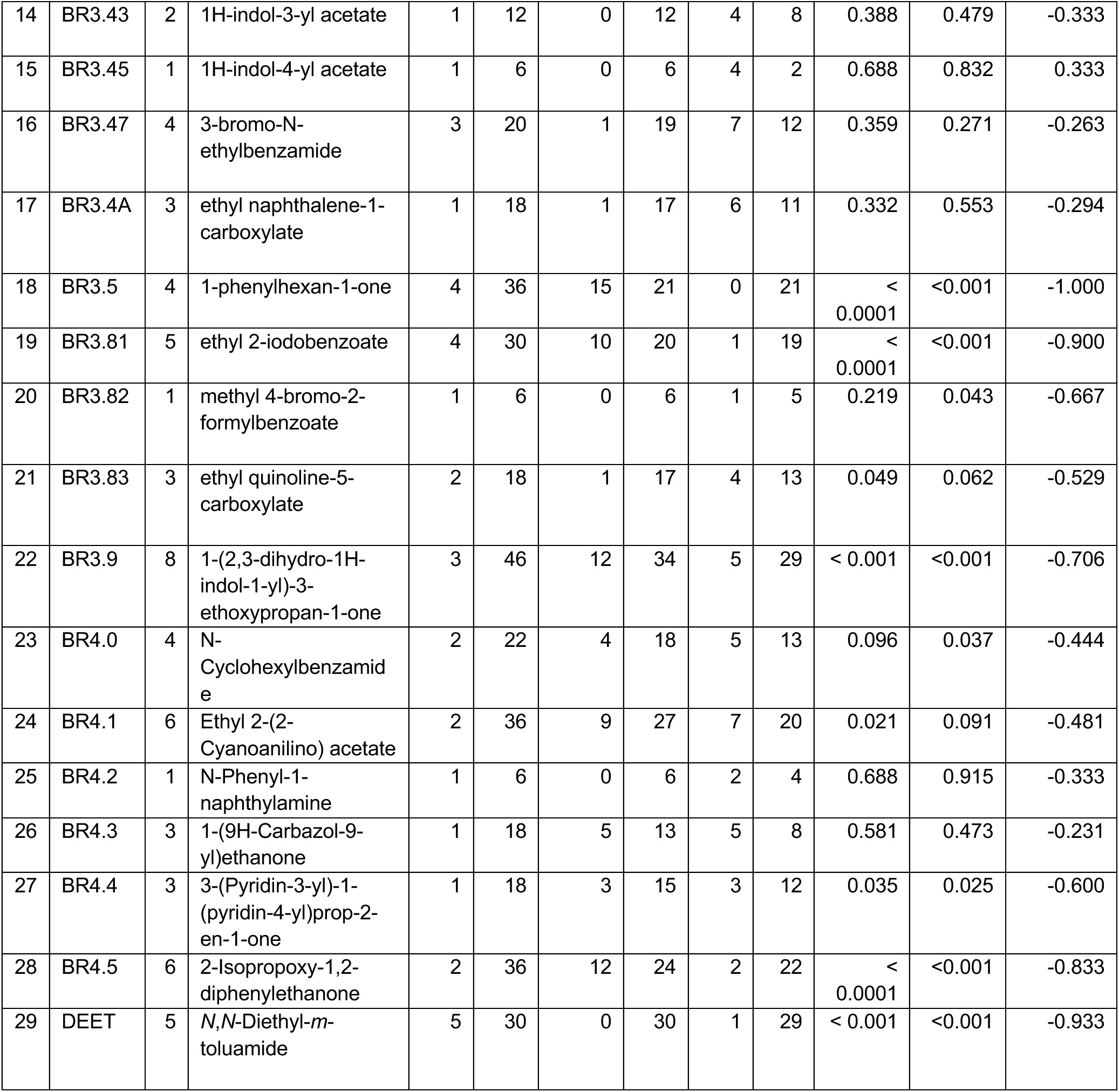
Repellency testing in the 2-choice plate assay from second iteratively trained round of predictions.

| # | Repellent Candidate | Batches Tested | Names of compounds | Colonies Tested | Plates Total | Not Participating | Participating | 1<sup>st</sup> Choice Repellent | 1<sup>st</sup> Choice Control | 1<sup>st</sup> Choice Control vs Repellent Mc Namar P | Honey Consumption Wilcoxon P | 1<sup>st</sup> Choice Preference Index* |
| --- | --- | --- | --- | --- | --- | --- | --- | --- | --- | --- | --- | --- |
| 1 | BR3.1 | 3 | phenyl 3-methoxypropanoate | 2 | 24 | 5 | 19 | 1 | 18 | < 0.0001 | <0.001 | -0.895 |
| 2 | BR3.10 | 1 | (1R,2R)-2-ethoxycyclohexan-1-ol | 1 | 6 | 0 | 6 | 1 | 5 | 0.219 | 0.344 | -0.667 |
| 3 | BR3.12 | 2 | 1-methyl-2,3-dihydro-1H-indole | 2 | 12 | 1 | 11 | 2 | 9 | 0.065 | 0.005 | -0.636 |
| 4 | BR3.15 | 3 | ethyl 2-(2-methylphenyl)acetate | 3 | 18 | 0 | 18 | 0 | 18 | < 0.0001 | <0.001 | -1.000 |
| 5 | BR3.1A | 1 | 2-Methyl-N-propylbenzamide | 1 | 6 | 3 | 3 | 2 | 1 | 1.000 | 0.180 | 0.333 |
| 6 | BR3.2A | 2 | 2-(3-Oxo-3-phenylpropyl)cyclopentanone | 2 | 11 | 0 | 11 | 4 | 7 | 0.549 | 0.127 | -0.273 |
| 7 | BR3.30 | 5 | N-(3-ethoxypropyl)-2-iodobenzamide | 3 | 29 | 3 | 26 | 4 | 22 | < 0.001 | <0.001 | -0.692 |
| 8 | BR3.35 | 1 | butyl 3-methylquinoline-4-carboxylate | 1 | 6 | 0 | 6 | 2 | 4 | 0.688 | 0.705 | -0.333 |
| 9 | BR3.38 | 1 | 3-bromo-N-propylbenzamide | 1 | 6 | 0 | 6 | 2 | 4 | 0.688 | 0.673 | -0.333 |
| 10 | BR3.3A | 6 | ethyl 2-(2,3-dihydro-1H-inden-1-yl)acetate | 5 | 36 | 15 | 21 | 0 | 21 | < 0.0001 | <0.001 | -1.000 |
| 11 | BR3.4 | 3 | 1-phenylhexan-3-one | 3 | 18 | 0 | 18 | 0 | 18 | < 0.0001 | <0.001 | -1.000 |
| 12 | BR3.40 | 2 | 2-butyl-2,3-dihydro-1H-indazol-3-one | 1 | 6 | 0 | 6 | 3 | 3 | 1.000 | 0.674 | 0.000 |
| 13 | BR3.42 | 5 | 1-{2-[(1E)-2-phenylethenyl]phenyl}ethan-1-one | 3 | 30 | 6 | 24 | 4 | 20 | 0.002 | 0.002 | -0.667 |
| 14 | BR3.43 | 2 | 1H-indol-3-yl acetate | 1 | 12 | 0 | 12 | 4 | 8 | 0.388 | 0.479 | -0.333 |
| 15 | BR3.45 | 1 | 1H-indol-4-yl acetate | 1 | 6 | 0 | 6 | 4 | 2 | 0.688 | 0.832 | 0.333 |
| 16 | BR3.47 | 4 | 3-bromo-N-ethylbenzamide | 3 | 20 | 1 | 19 | 7 | 12 | 0.359 | 0.271 | -0.263 |
| 17 | BR3.4A | 3 | ethyl naphthalene-1-carboxylate | 1 | 18 | 1 | 17 | 6 | 11 | 0.332 | 0.553 | -0.294 |
| 18 | BR3.5 | 4 | 1-phenylhexan-1-one | 4 | 36 | 15 | 21 | 0 | 21 | < 0.0001 | <0.001 | -1.000 |
| 19 | BR3.81 | 5 | ethyl 2-iodobenzoate | 4 | 30 | 10 | 20 | 1 | 19 | < 0.0001 | <0.001 | -0.900 |
| 20 | BR3.82 | 1 | methyl 4-bromo-2-formylbenzoate | 1 | 6 | 0 | 6 | 1 | 5 | 0.219 | 0.043 | -0.667 |
| 21 | BR3.83 | 3 | ethyl quinoline-5-carboxylate | 2 | 18 | 1 | 17 | 4 | 13 | 0.049 | 0.062 | -0.529 |
| 22 | BR3.9 | 8 | 1-(2,3-dihydro-1H-indol-1-yl)-3-ethoxypropan-1-one | 3 | 46 | 12 | 34 | 5 | 29 | < 0.001 | <0.001 | -0.706 |
| 23 | BR4.0 | 4 | N-Cyclohexylbenzamide | 2 | 22 | 4 | 18 | 5 | 13 | 0.096 | 0.037 | -0.444 |
| 24 | BR4.1 | 6 | Ethyl 2-(2-Cyanoanilino) acetate | 2 | 36 | 9 | 27 | 7 | 20 | 0.021 | 0.091 | -0.481 |
| 25 | BR4.2 | 1 | N-Phenyl-1-naphthylamine | 1 | 6 | 0 | 6 | 2 | 4 | 0.688 | 0.915 | -0.333 |
| 26 | BR4.3 | 3 | 1-(9H-Carbazol-9-yl)ethanone | 1 | 18 | 5 | 13 | 5 | 8 | 0.581 | 0.473 | -0.231 |
| 27 | BR4.4 | 3 | 3-(Pyridin-3-yl)-1-(pyridin-4-yl)prop-2-en-1-one | 1 | 18 | 3 | 15 | 3 | 12 | 0.035 | 0.025 | -0.600 |
| 28 | BR4.5 | 6 | 2-Isopropoxy-1,2-diphenylethanone | 2 | 36 | 12 | 24 | 2 | 22 | < 0.0001 | <0.001 | -0.833 |
| 29 | DEET | 5 | N,N-Diethyl-m-toluamide | 5 | 30 | 0 | 30 | 1 | 29 | < 0.001 | <0.001 | -0.933 |

**Table S3:** Initial Data for Round 1 Machine Learning.

| PubChem CID | Repellency |
| --- | --- |
| 102911 | Repel |
| 102981 | Repel |
| 1032 | Repel |
| 10402 | Inactive |
| 10931 | Inactive |
| 10976 | Repel |
| 111037 | Inactive |
| 1127 | Repel |
| 11357 | Repel |
| 11369 | Inactive |
| 11419 | Repel |
| 11579 | Inactive |
| 12291 | Repel |
| 12310947 | Inactive |
| 123233 | Inactive |
| 12403 | Repel |
| 12404 | Repel |
| 1254 | Inactive |
| 12587 | Inactive |
| 12901 | Inactive |
| 13002 | Inactive |
| 13008 | Repel |
| 13566 | Repel |
| 13629102 | Inactive |
| 136888 | Repel |
| 137414 | Repel |
| 13879 | Repel |
| 14240 | Inactive |
| 15105174 | Inactive |
| 15133 | Inactive |
| 1549026 | Inactive |
| 15624 | Repel |
| 16678 | Inactive |
| 170833 | Repel |
| 17100 | Inactive |
| 17355 | Repel |
| 176 | Repel |
| 180 | Inactive |
| 18954 | Inactive |
| 19079 | Inactive |
| 199294 | Inactive |
| 20745 | Inactive |
| 221558 | Repel |
| 225147 | Repel |
| 232703 | Inactive |
| 233220 | Inactive |
| 2345 | Repel |
| 235640 | Inactive |
| 240 | Repel |
| 244 | Repel |
| 24623 | Repel |
| 246831 | Repel |
| 246835 | Repel |
| 24847 | Inactive |
| 24981 | Repel |
| 24994 | Repel |
| 261 | Repel |
| 2879 | Inactive |
| 2969 | Inactive |
| 3004112 | Inactive |
| 3026 | Repel |
| 3102 | Repel |
| 311 | Inactive |
| 31229 | Repel |
| 31263 | Repel |
| 314154 | Inactive |
| 33191 | Repel |
| 3438 | Repel |
| 346056 | Inactive |
| 4004 | Inactive |
| 4115 | Inactive |
| 4133 | Repel |
| 428 | Repel |
| 4284 | Inactive |
| 442484 | Inactive |
| 454 | Repel |
| 522168 | Repel |
| 527 | Repel |
| 5282108 | Inactive |
| 5284503 | Inactive |
| 5312738 | Repel |
| 53167 | Repel |
| 532460 | Repel |
| 5354264 | Repel |
| 5354896 | Inactive |
| 5355850 | Inactive |
| 5356959 | Inactive |
| 5363140 | Repel |
| 5365800 | Repel |
| 5370101 | Repel |
| 5462719 | Inactive |
| 5634 | Inactive |
| 6028 | Inactive |
| 61004 | Inactive |
| 61067 | Repel |
| 61417 | Repel |
| 62350 | Inactive |
| 637511 | Repel |
| 637564 | Inactive |
| 637566 | Inactive |
| 638011 | Inactive |
| 638014 | Inactive |
| 64685 | Inactive |
| 6551 | Inactive |
| 6569 | Repel |
| 6590 | Repel |
| 66093 | Repel |
| 6673 | Inactive |
| 6781 | Repel |
| 6915 | Inactive |
| 6923 | Inactive |
| 69389 | Inactive |
| 69656 | Repel |
| 69702 | Inactive |
| 6985 | Repel |
| 6989 | Inactive |
| 70316 | Repel |
| 712 | Inactive |
| 7298 | Repel |
| 7410 | Repel |
| 7416 | Repel |
| 74741 | Repel |
| 7497 | Repel |
| 7507 | Inactive |
| 75200 | Inactive |
| 75454 | Inactive |
| 7554 | Repel |
| 75678 | Repel |
| 7590 | Repel |
| 75945 | Inactive |
| 7618 | Inactive |
| 76194 | Repel |
| 76388 | Inactive |
| 7677 | Inactive |
| 7678 | Repel |
| 7719 | Repel |
| 77321 | Inactive |
| 77323 | Inactive |
| 7746 | Repel |
| 7751 | Repel |
| 7757 | Repel |
| 7761 | Repel |
| 7762 | Repel |
| 7777 | Inactive |
| 77844 | Inactive |
| 7794 | Inactive |
| 78481 | Inactive |
| 7854 | Repel |
| 7912 | Repel |
| 7923 | Repel |
| 79306 | Repel |
| 7965 | Repel |
| 8007 | Repel |
| 8050 | Inactive |
| 8051 | Repel |
| 8094 | Inactive |
| 8096 | Inactive |
| 8102 | Repel |
| 81188 | Repel |
| 8148 | Repel |
| 8149 | Inactive |
| 8180 | Inactive |
| 8222 | Repel |
| 8257 | Inactive |
| 8268 | Inactive |
| 8294 | Inactive |
| 8332 | Inactive |
| 8471 | Repel |
| 8554 | Repel |
| 86079 | Inactive |
| 8635 | Inactive |
| 8797 | Repel |
| 8815 | Inactive |
| 8883 | Inactive |
| 8902 | Repel |
| 8916 | Repel |
| 95315 | Inactive |
| 95500 | Repel |
| 95633 | Repel |
| 969491 | Inactive |
| 98279 | Repel |
| 9862 | Repel |
| 996 | Repel |
| 999 | Inactive |

**Table S4:** Data added for Round 2 after testing for Bee Repellency.

| PubChem CID | Repellency |
| --- | --- |
| 4284 | Repel |
| 6877 | Repel |
| 82336 | Repel |
| 6660 | Repel |
| 20745 | Inactive |
| 7802 | Repel |
| 7288 | Inactive |
| 76398 | Inactive |
| 77224 | Inactive |
| 10953 | Inactive |
| 14286 | Inactive |
| 60993 | Repel |
| 98218 | Inactive |
| 12665 | Inactive |
| 12753 | Inactive |
| 69326 | Inactive |
| 61265 | Inactive |
| 7410 | Inactive |
| 347476 | Repel |

